# Chemotherapy dose scheduling via Q-learning in a Markov tumor model

**DOI:** 10.1101/2025.06.05.658118

**Authors:** M. Giles, P.K. Newton

## Abstract

We describe a Q-learning approach to optimized chemotherapy dose scheduling in a stochastic finite-cell Markov process that models tumor cell natural selection dynamics. The three competing subpopulations comprising our virtual tumor are a chemo-sensitive population (S), and two chemo-resistant populations, *R*_1_ and *R*_2_, each resistant to one of two drugs, *C*_1_ and *C*_2_. The two drugs are toggled off or on which constitute the actions (selection pressure) imposed on our state-variables (*S, R*_1_, *R*_2_), measured as proportions in our finite state-space of *N* cancer cells (*S* + *R*_1_ + *R*_2_ = *N*). After the converged chemo-dosing policies are obtained, corresponding to a given reward structure, we focus on three important aspects of chemotherapy dose scheduling. First, we identify the most likely evolutionary paths of the tumor cell populations in response to the optimized (converged) policies. Second, we quantify the robustness in the ability to reach our target of balanced co-existence in light of incomplete information in both the initial cell-populations as well as the state-variables at each step. Third, we evaluate the efficacy of simplified policies which exploit the symmetries uncovered from an examination of the full policy. Our reward structure is designed to delay the onset of chemo-resistance in the tumor by rewarding a well-balanced mix of co-existing states, while punishing unbalanced subpopulations to avoid extinction.

## I. INTRODUCTION

The optimal design of multi-drug chemotherapy scheduling is a combinatorially daunting task given the many possible combinations of chemotherapy drugs available [1] and the multitude of mechanisms by which they can act on heterogenous tumor cell populations with a broad range of mutational profiles [2, 3]. Because of these complexities and the high-stakes consequences associated with decision making, most chemotherapy scheduling follows tried and true, time-tested protocols, such as maximum tolerated dosing (MTD) or metronomic dosing [4], using small and predictable combinations of drugs comprising the chemotherapy cocktails. The benefit of using pre-scheduled standard regimens are the years of experience that have been invested in understanding the dose-response curves under different scenarios [4]. Because of this deep well of experience, results are fairly predictable and considered safe over a wide range of patient populations. Despite this, very few oncologists would argue that the pre-scheduled (fixed) protocols currently in use are adaptively optimized to treat cell populations that are constantly in flux due to selection pressure from a multitude of sources [5]. In fact, it is known that many of the MTD schedules can actually select for resistant tumor-cell populations (by killing off the sensitive population), thereby enhancing the possibility of recurrance due to developed resistance. An eco-evolutionary analogy has been documented both in insect/pest populations and microbial populations where repeated heavy use of a single toxin (DDT, antibiotics) kills the sensitive cells, releasing the resistant cells from competition with their higher-fitness sensitive competitors, allowing them to take over the ecological niche, a process known as competitive release [5–8].

One might then wonder what an optimized dosing schedule would look like if designed by a computational reinforcement learning algorithm using repeated trials that accumulate rewards for success, and punishments for failure, that attempts to steer the competitive balance among the different cell populations towards some beneficial target state. The potential power of developing an algorithmic approach is that it can, in principle, be used to design much more complex protocols using many more drug combinations in a higher-dimensional setting with a more fine-grained delineation of subpopulations. To gain an appreciation for how daunting the task is of designing multi-drug cocktails that target different evolving subpopulations, consider a simple design of two-drug cocktails from a choice of ten possible chemotoxins — this requires the evaluation of dose-response curves for 45 different combinations (ten choose two), all in a dynamic and stochastic environment in which the cocktails can be remixed and administered temporally. Drug cocktails that make use of four drugs from a choice of ten require the evaluation of 210 different combinations. However there are hundreds of drugs currently in use that target cancer cell populations with different mutational profiles that determine a wide range of proliferation rates, and combining them in four or more combinations is not unusual. All of these mixtures could (and should) change dynamically as the balance of cell types changes, which leads to a daunting combinatorial and dynamical decision making process of adaptive stochastic optimization and nonlinear control [1].

This paper is not the first to propose the use of reinforcement learning as a control theory method applied to evolving populations. References [9–11] all provide nice comprehensive reviews of reinforcement learning approaches in the context of precision oncology, while [12] provides a general review of computational approaches in oncology. Ref [13] provides an example of a model-free RL method introduced for closed-loop control of chemotherapy drug-dosing based on a three-state deterministic model, along with a chemotherapy control function. Ref. [14] develops a RL approach to learning effective drug-cycling policies to fight anti-microbial resistance in an evolving context, while [15] develops an optimization evolutionary game theory model for drug-dosing strategies.

Our goal in this paper is to implement a reinforcement learning algorithm in a relatively low-dimensional setting to design optimal schedules for a two-drug chemotherapy protocol, with three subpopulations of competing tumor cells with differing responses to each drug. We will discuss the advantages and opportunities associated with using algorithmically designed dosing schedules in the final section.

## II. CELL POPULATION DYNAMICS OF THE TUMOR MODEL

### A. Finite-cell discrete-time evolutionary game

In this work, cancer evolution is modeled as a discrete time, stochastic evolutionary game [8]. It takes the form of a birth-death process between three competing cancerous cell subpopulations: *S*, the universally sensitive cell type; *R*_1_, resistant to *C*_2_ but sensitive to *C*_1_; *R*_2_, resistant to *C*_1_ but sensitive to *C*_2_. Here, *C*_1_ and *C*_2_ represent two different chemotoxins that are effective against the sensitive cell type *S*, but to each of which one of the cell types *R*_1_ or *R*_2_ expresses a phenotypic resistance.

The system state is given by the vector 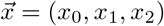, where *x*_0_, *x*_1_, and *x*_2_ are the population of *S, R*_1_, and *R*_2_ cells respectively. The total population given by *x*_0_ + *x*_1_ + *x*_2_ is fixed and denoted *N*. Thus the state space for this system is given by 𝕊_*N*_ = {(*x*_0_, *x*_1_, *x*_2_) *ϵ* ℕ^3^ : *x*_0_ + *x*_1_ + *x*_2_ = *N*}. For ease of notation, *S, R*_1_, and *R*_2_ type cells will sometimes be called 0-, 1-, or 2-type cells, or *i*-type in the general case. The state space 𝕊_*N*_ can then be represented as a triangular simplex in two dimensions, where vertices represent the points 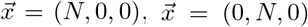, and 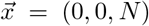. Faces of the triangular simplex (shown in figure 1) represent the points where one species is extinct, (i.e. *x*_*i*_ = 0, *i* = 1, 2 or 3), and the interior of the simplex represents states where all cell types co-exist. An example of a discrete grid for *N* = 50 is shown in figure 1. The midpoint of the triangular simplex is the state of maximal co-existence (*S, R*_1_, *R*_2_) = (*N/*3, *N/*3, *N/*3). Since this point is the furthest from all three boundaries, in a stochastic environment, it is the least likely position for stochastic fluctuations to cause extinction.

**FIG. 1.**
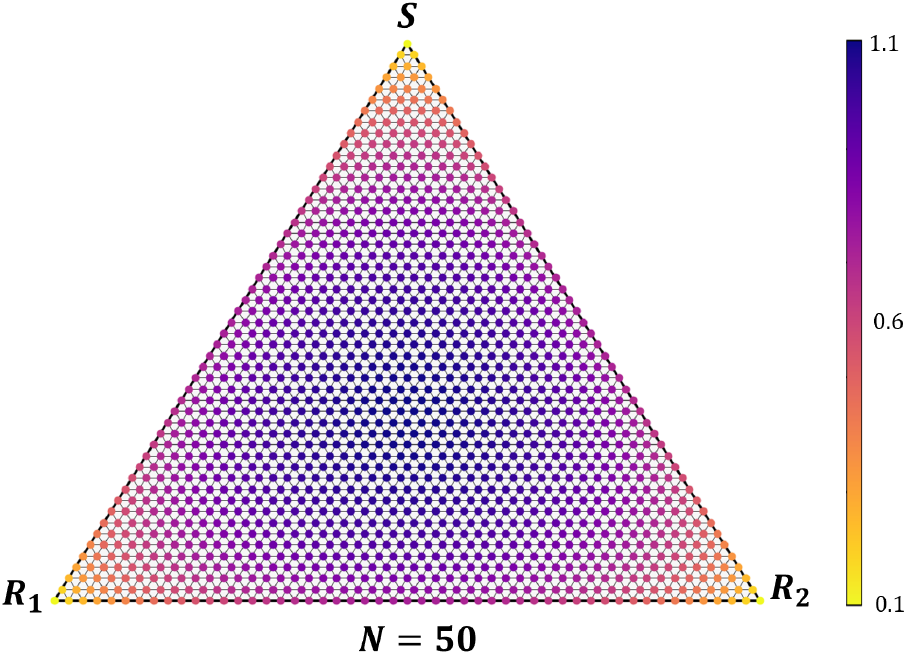
Entropy heat map (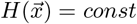. color bar on right) on the triangular simplex (*S* + *R*_1_ + *R*_2_ = *N*) describing the discrete state space 𝕊_50_.

To quantify interactions between the three cell subpopulations, a payoff matrix, *M*, is given by

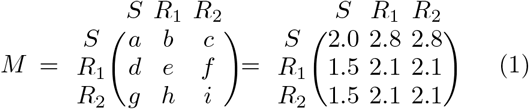

Entries of this matrix correspond to the assumed reproductive reward a cell receives at each time step, based on the other cells it encounters. Row labels reflect the cell type receiving the reward, and columns reflect its competitor. The top row dominates the two lower rows, reflecting the fitness cost associated with resistance, well documented in microbial and other competing populations [16]. Because of this gap in the fitness between *S* and *R*_1,2_ the *S* population will ultimately reach fixation in the absence of chemotherapy (it represents a Nash equilibrium of the system). In our model we take both resistant populations to have symmetric fitness values, but are resistant to two different drugs *C*_1_ and *C*_2_. At the same time, two interacting cells of a non-dominant type receive greater payoff than two interacting *S* type cells. Thus the game naturally encourages evolution of *S* type cells, even though a system saturated with *S* cells receives less total payoff than a population saturated with resistant cells. This provides a good model for the scenario under investigation, since the *S* population is expected to outcompete resistant cells in the absence of chemotherapy, but a fully sensitive population will be more vulnerable to chemotherapy providing a selective advantage to the resistant population.

It is an interesting fact that the structure of this payoff matrix corresponds to a prisoner’s dilemma (PD) game between *S* and *R*_1_ and between *S* and *R*_2_. That is, *b* > *e* > *a* > *d*, and *c* > *i* > *a* > *g*. A PD is a domination-class game, where the greatest payoff is always received by one dominant strategy (cell type), in this case *S*. This is due to the *fitness cost of resistance* discussed in [16] for microbial populations, and [17] for tumor cells. In addition to these PD games, a neutral game is established between *R*_1_ and *R*_2_, defined by the relation *e* = *f* = *h* = *i*, to impose a symmetry relation among these species.

It is assumed that every cell in the system has an equal probability of interacting with every other cell (i.e the well-mixed assumption). Given the payoff matrix *M* and population state 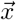, the expected payoff *F*_*i*_ received by a cell of type *i* is:

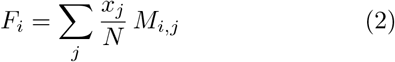

where *x*_*j*_*/N* is the probability that the focal cell interacts with a *j*-type cell and *M*_*i,j*_ is the *i, j*^*th*^ entry of *M*, corresponding to the reward the cell would receive against a cell of type *j*. In the absence of chemotherapy, *F*_1_ *> F*_2_ = *F*_3_ for all three states corresponding to the vertices of the state-space triangle.

Survival and reproduction prowess is not solely governed by the payoff matrix, but also by the evolving proportions of each of the cell types in the population. To capture the intensity of selection in this model (which we will use to apply chemotherapy dosing), a parameter 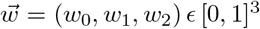 is introduced, with elements *w*_*i*_ corresponding to the selection pressure imposed on cells of type *i*. Fitness *f*_*i*_ of a cell of type *i* is then defined as:

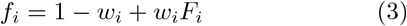

Thus as *w*_*i*_ → 1 (strong selection limit), *f*_*i*_ → *F*_*i*_ and selection strongly governs evolution. On the other hand, as *w*_*i*_ → 0 (weak selection limit), selection becomes nonexistent and the evolutionary dynamics of the system approach neutral drift. Given that chemotherapy imposes selection pressure on different cell sub-types, we use the selection parameter as our chemotherapy control knob described in the next section.

### B. The Markov process

The birth-death process then proceeds on a discrete grid as shown in Figure 1. At each time step, one cell is chosen for elimination (e) and another for reproduction (r). Both choices are random and independent; the cell to be eliminated is chosen with uniform probability, while the cell to reproduce is chosen with probability weighted by fitness. Specifically:

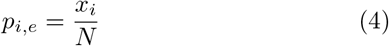

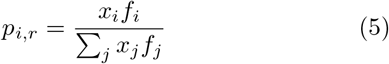

where *p*_*i,e*_ and *p*_*i,r*_ are the probabilities that a cell of type *i* is chosen for elimination or reproduction, respectively. Once the cells have been chosen, the population of the eliminated cell type decreases by one, and that of the reproducing cell type increases, while the total population remains fixed at *N*. In the case that the eliminated and reproducing cells are of the same type, the system state does not change. The stochastic process describes a weighted Moran process [8] that can be thought of as an evolutionary game [18] which, in the absence of chemotherapy will result in the *S* population reaching fixation.

In this model there is no mechanism for mutation, and so by equation 5, if the cell type *i* becomes extinct (i.e. *x*_*i*_ = 0) then *p*_*i,r*_ = 0 and the species can never reemerge (i.e. each boundary is an absorbing state of the system). Similarly, if the cell type *j* has saturated the system (i.e. *x*_*j*_ = *N, x*_*i*_ = 0 ∀ *i* ≠ *j*), then by equations 4 and 5, *p*_*j,e*_ = *p*_*j,r*_ = 1, and the system has also reaches an absorbing state. Viewing S_*N*_ as a triangular simplex, extinction of one species corresponds to an edge of the simplex, while extinction of two corresponds to a vertex, and both are inescapable.

Now, define 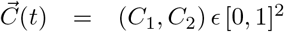 to be the chemotherapy dose administered to the system, where the elements *C*_1_ and *C*_2_ correspond to the doses of drugs *C*_1_ and *C*_2_. Time *t* in our model is discrete. To introduce chemotherapy into the model, we use our selection pressure parameter associated with how each of the phenotypes responds to each of the drugs, defined in eqn (3). Let 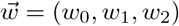, so that chemotoxin concentrations can alter the selection pressure imposed on each cell type:

- w0 = 1™min(1, C1 + C2)
- w1 = 1 ™ C2
- w2 = 1 ™ C1

This formulation of 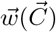 encodes the resistance profile of each of the cell types *S, R*_1_, and *R*_2_, so that chemotoxin concentration adversely affects the selection pressure, and therefore fitness (by eqn (3)), of only non-resistant cell species.

## III. REINFORCEMENT LEARNING

### A. A Primer on Q-Learning

Reinforcement Learning (RL) is particularly well-suited to control optimization in discrete-time stochastic processes [19]. An RL agent can sense the state *s* of its environment and choose a corresponding action *a* to take. The state is then updated according to some transition function **T**(*s, s*^*′*^, *a*), where *s*^*′*^ is the updated state, and the agent is rewarded or punished according to a reward function **R**(*s, s*^*′*^, *a*). The agent uses this reward to assign a value to its chosen action. Through repeated trial and error, the agent refines the value assigned to each action in the action space (𝔸), at every state in the state space 𝕊, to develop (i.e. “learn”) an optimal control policy *π* for maximizing its expected reward. Thus a reinforcement learning agent must operate on a Markov Decision Process (MDP), defined by the 4-tuple (𝕊, 𝔸, **T, R**). In the context of this work, 𝕊 is the set of valid subpopulation distributions (*x*_0_, *x*_1_, *x*_2_), and 𝔸 is the set of allowable chemotherapy doses. **T** is then described by birth-death probabilities per the model formulated in Section II B, and **R**(*s, s*^*′*^, *a*) = **R**(*s*^*′*^) is a table assigning values to the state reached by the agent over the previous transition step (see Section III B). These algorithms typically involve an initial exploration phase where the agent performs this trial and error learning sequence, followed by an exploitation phase, where the agent uses the policy it has learned to maximize the benefit it receives from its environment.

RL techniques can be subdivided into model-based and model-free techniques [19]. When the process being investigated evolves according to a known model, there exist (model-based) RL methods that can leverage this information to quickly learn effective policies. When no such model is available, model-free techniques must be implemented to learn new policies from experimental data. In this work we develop a model-free approach.

Q-learning is one such model-free technique. It esimates a quality function, **Q**(*s, a*), describing the expected reward for taking the action *a*, given that the system is at state *s*. An optimal policy can then be extracted as *π*(*s*) = argmax_*a*_**Q**(*s, a*). The pseudocode provided in Algorithm 1 provides a fairly standard implementation of synchronous Q-learning.

#### Algorithm 1 Synchronous Q-Learning

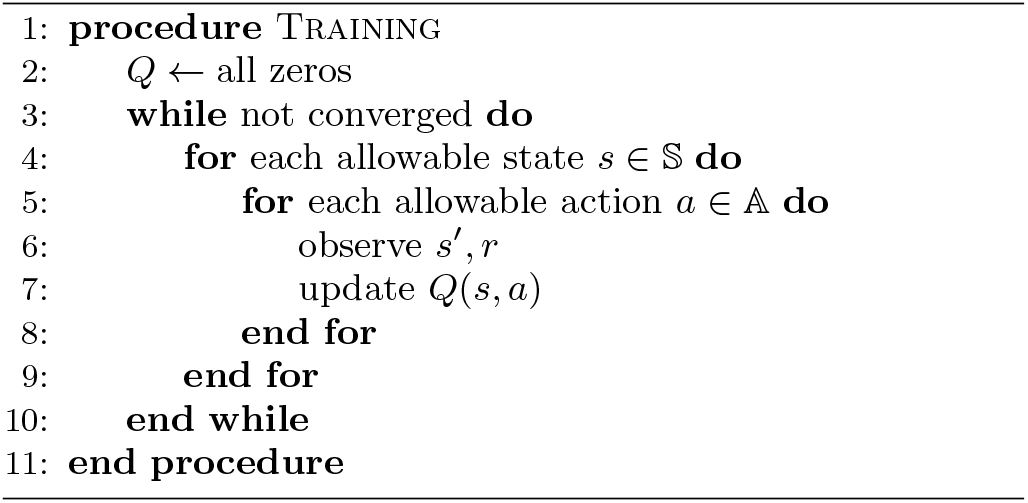

At each step of the inner for loop, *s*^*′*^ is the state that the system transitions to as a result of taking action *a* from state *s*. The reward received for achieving this state *s*^*′*^ is given by *r*. The transition from *s* to *s*^*′*^ is simulated using the weighted Moran process provided in section II. The RL agent explores the state-space environment, attempting each action *a* at every state *s* and observing only the resultant new state *s*^*′*^. Through repeated attempts and observation, the agent may infer transition probabilities for the various allowable state-action pairs, and in this way computes the quality function. After observing the new state *s*^*′*^ and receiving the reward *r* associated with this new state, the update equation for *Q*(*s, a*) is given by the Bellman equation [19]:

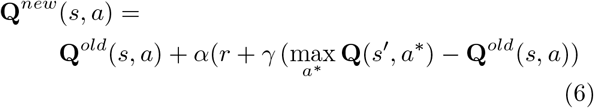

The parameters *α* and *γ* can be adjusted to tailor the learning process. *α* represents the learning rate, which governs how quickly the RL agent is able to adapt its estimate of **Q** in response to experimental trials. *δ ϵ* (0, 1] is a discount factor which devalues future rewards relative to immediate rewards, motivating the agent to reach desirable states quickly.

### B. The reward structure

Per the payoff matrix in eqn (1), *S* cells outcompete *R*_1_ and *R*_2_ cells in the absence of chemotherapy and the *S* population will reach fixation. By contrast, administering chemotherapy with high doses can eliminate the *S* population, enabling the resistant populations to invade the system leaving the patient with a resistant tumor. Our goal is to apply chemotherapy in such a way that all three subpopulations *S, R*_1_, and *R*_2_ co-exist, competing on an equal footing without allowing any to reach fixation.

The model presented in Section II is stochastic and therefore features an element of random drift [18]. The smaller the population of any cell species, the more likely it can become extinct due to stochastic fluctuations. It is therefore advantageous to achieve and maintain maximal species diversity in order to avoid extinction by random drift. Quantitatively, this is captured by a maximization of entropy at the equal co-existence state given by:

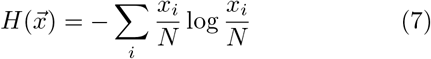

This notion of entropy is that of Shannon entropy in information theory, [20, 21] and has more recently demonstrated relevance in describing tumor complexity and heterogeneity [22][23][24]. A plot of the entropy for the simplex representing 𝕊_50_ is provided in Figure 1

To encourage the Q-learning agent to maximize species diversity, it should be strongly rewarded for achieving high-entropy states and punished (i.e. assigned a negative reward) for reaching low-entropy states (near the boundaries). The reward structure we use, based on piecewise constant entropy values, is shown in Figure 2. The entropy-based reward structure assigns a strong reward (+1) to the states that achieve an entropy value within 0.05% of the maximum value, and a punishment (reward of −1) to states near the boundaries. The region assigned a reward of (+1) is our *target* region. The structures shown in Figures 2b and 2c add small, widening buffer regions to the target, in an attempt to guide the RL agent toward it. To evaluate the importance of reward structure, chemotherapy policies were trained using the three piecewise constant entropy based reward structures and the results compared.

### C. Implementation

In the context of this work, the state space 𝕊_*N*_ is the set of allowable states 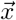, each corresponding to a unique sub-population balance among the three cell species discussed in Section II A. The action space 𝔸 is defined as the space of allowable chemotherapy concentrations 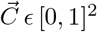 (see Section II B). To minimize computational load associated with learning the quality function *Q*, the action space 𝔸 := {(0, 0), (0, 1), (1, 0)} was selected, for a total of just three admissible actions. The first corresponds to both *C*_1_ and *C*_2_ turned off (no chemotherapy), the second has *C*_1_ turned off, while *C*_2_ is on, while the third has *C*_1_ on, while *C*_2_ is off. The action 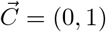 puts selection pressure on the *S* and *R*_2_ cells which are sensitive to the *C*_2_ chemotoxin. Similarly, 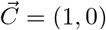 puts selection pressure on the *S* and *R*_1_ cells which are both sensitive to the *C*_1_ chemotoxin. 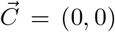 allows the *S* subpopulation to reach fixation since in the absence of chemotherapy, *S*-type cells receive greater payoff and therefore outcompete the resistant populations (see Equation 1). Thus this choice of 𝔸 features one action to favor each of the three cell subtypes *S, R*_1_, and *R*_2_, and was therefore determined to be the simplest action space that still provides sufficient agency in controlling the population’s evolutionary trajectory. For all analyses, a total population size of *N* = 100 was chosen. For values smaller than this, the state space 𝕊_*N*_ is too susceptible to random drift, and therefore difficult to consistently control. As *N* increases, the role of selection becomes more significant in governing population dynamics. At very large values (*N* → ∞), the evolutionary dynamics become deterministic (i.e. no drift). The value *N* = 100 provides a comfortable middle ground where stochastic phenomena are still observable but do not overwhelmingly dominate the dynamics. Importantly, this value for *N* was also sufficiently small to allow for reasonably quick computational convergence of the *Q* estimate.

For each of the reward structures shown in Figure 2, training proceeded according to Algorithm 1. Preliminary testing discovered that training parameter values of *α* = 0.05 and *δ* = 0.9 resulted in stable and relatively rapid convergence of successful chemotherapy policies. Note that, while absolute convergence of Q-learning is guaranteed in the limit of infinite training cycles [25][26], it was found through early experimentation that the incremental benefit of additional training time declines severely after the first few tens of thousands of training iterations. This is exemplified in Figure 3, which shows the absolute change in Q-value over all state-action pairs versus the number of training cycles.

In the interest of mitigating computational burden, training was terminated after 100, 000 iterations of the outer while loop, which was found to be sufficiently long for generating effective chemotherapy schedules (see section IV A).

## IV. RESULTS

### A. Learned policies

We turn now to a description of the results. Three chemotherapy policies were produced, each corresponding to one of the reward structures presented in Figure 2. The policies are visually represented by Figures 4a-4c. The policies prescribe chemotherapy doses (i.e. actions) based solely on the current system state. Formally, the policy is constructed by choosing, at each state 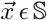, the action 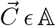 that maximizes the learned quality function, 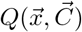.

**FIG. 2.**
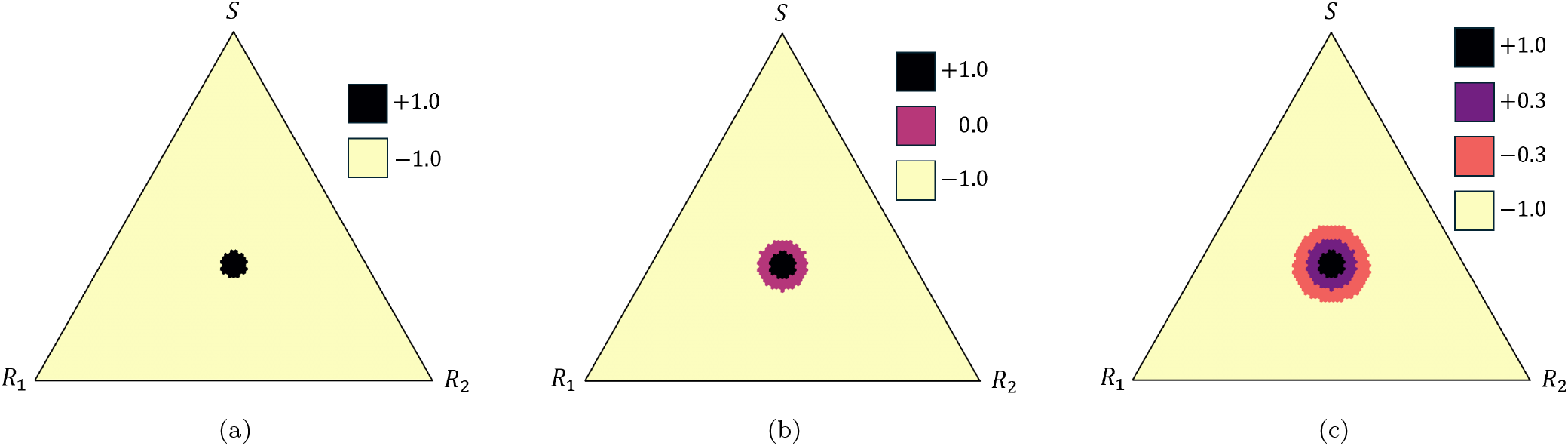
Reward structures based on piecewise constant approximations of the smooth entropy landscape. (a) Two-region reward structure; (b) Three-region reward structure; (c) Four-region reward structure.

**FIG. 3.**
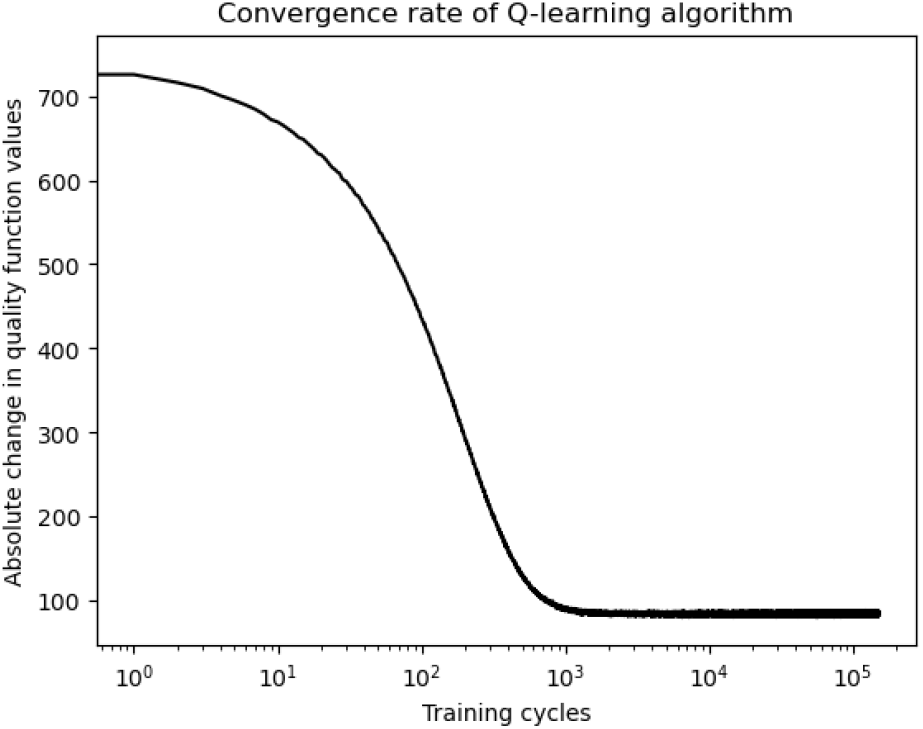
Net change in *Q* for all state-action pairs as a function of training cycles.

**FIG. 4.**
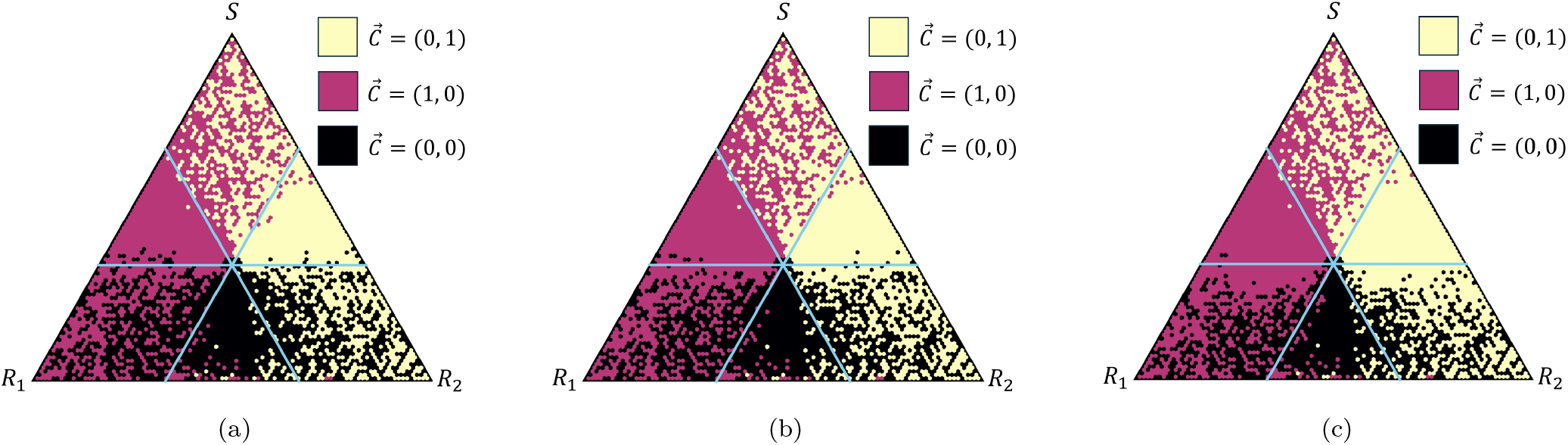
Three learned policies using the reward structures in figure 2. Symmetry lines divide the triangle into six distinct zones. The three triangular shaped zones favor a single policy that guides the trajectory towards the opposite corner. The three trapezoidal shaped zones favor a random mixture of the two adjoining triangular policies. (a) Two-region (fig 2(a)); (b) Three-region (fig 2(b)); (c) Four-region (fig 2(c)).

The first observation is that the three learned policies are qualitatively similar and exhibit an important symmetry. The triangular simplex on 𝕊 is divided into six distinct zones. The three corner zones prescribe a balance between two of the three available actions, while the zones opposing the corners each prescribe a single action. The policies shown in Figures 4a-4c have dividing lines drawn across the *N/*3 axis for each subpopulation, which helps to distinguish these six zones. Recall that the three allowable actions each favor population growth for one of the three cell species. In the three zones opposing each corner, only one of the subpopulations is at risk of extinction (i.e. a trajectory hitting the side-edge). As a result, the prescribed action is the one that favors the at-risk cell type. On the other hand, in each corner zone there are two subpopulations at risk of extinction (i.e. a trajectory hitting a corner).The resulting policy strikes a balance between the actions favoring each of them. While the learned policies shown in Figures 4b-4c have the same general six-zone structure, they differ slightly near the bounds of each zone. Our general conclusion, however, is that since the different reward structures from Figure 2 does not much impact the chemotherapy policy produced, these policies are relatively robust to changes in the reward structure that build in finer gradients to the entropy landscape. The following sections will investigate the general performance of these six-zone policy structures.

### B. Most likely paths

To evaluate the effectiveness of a learned policy, we compute the locally most probable state trajectories (paths) that the policy yields for six initial conditions, one in each symmetry zone. This provides insight into the general behavior that the policy attempts to elicit from the system. To compute this path, at each step, we choose the neighboring site with the largest transition probability.

More generally, given some initial condition 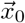 and a predetermined chemotherapy schedule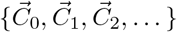, the most likely state trajectory from 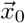 can be infor-mally described by the sequence 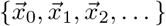, where for each time-step *t* ≥ 1,

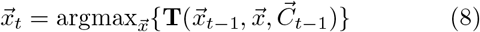

That is, at each time step we simply choose the state that the system is most likely to transition to, which will depend only on the previous time step’s state and chemotherapy dosing. We call this the *locally* most probable trajectory as it only seeks to maximize one-step transition probabilities between states.

To evaluate learned policies, the chemotherapy schedule 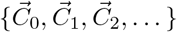 in the procedure above is chosen according to the policy’s suggestions. As described in Section III A, a policy *π* is a mapping between from the state space to the action space (*π* : **S** → **A**). Thus to evaluate *π*, choose 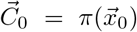. Transition probabilities can then be computed for the first transition (take 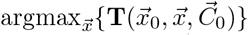) to find the most likely candidate for 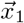. Then, 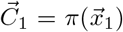, enabling computation of 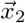, etc. In this way, the (locally) most probable state-trajectory and chemotherapy schedule corresponding to a learned policy and initial condition can be computed concurrently.

As a final technical detail, when computing each new element of the state-trajectory, previous states visited along the trajectory are removed from consideration. The reasoning for this decision is as follows: suppose the sequence 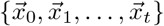 has been computed and that from state 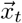 the most likely transition for the system is to remain at the same state. Then 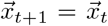 and so 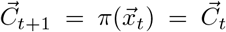. Therefore, nothing has changed in the system, so in the next time step, the system will once again be most likely to remain in the same state. The result is that the sequence gets stuck, with 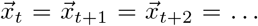 for all time (i.e. a probabilistic fixed point). A similar behavior would also be observed if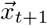 was taken to be any other previously visited state 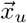 (for 0 ≤*u < t*), except the trajectory would get stuck in a cycle of states rather than a single state. However, in any realization of the underlying process, stochasticity would eventually push the state out of any such loop (provided the system had not reached one of its three fixed points). Thus these cycles are not important features of the learned chemotherapy schedules, and are avoided when computing locally most likely trajectories. Equa-tion 8 can then be better written as the system:

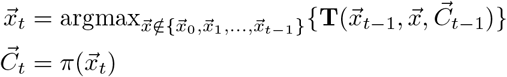

for a given initial condition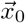 and policy *π*.

Figure 5 shows the most likely paths taken from each of the six initial conditions 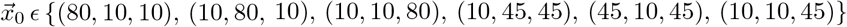. The initial conditions (ICs) were chosen to provide a representitive set from each zone. Each of the six computed trajectories reaches the target region, which is defined as the region where, per the reward structures in Figure 2, the RL agent receives the maximum reward of 1. The policies produced by the three different reward structures all performed very similarly. Considering the qualitative similarities in their structure, this does not come as a surprise.

To verify these results, stochastic simulations were performed from each of the initial conditions. Details on the implementation of these trials are introduced in Section IV E. From each initial condition, a set of 1000 random trajectories was simulated using transition probabilities described by the model presented in Section II B, conditioned on chemotherapy actions being taken according to the two-region policy. From these 1000 trajectories, a probability of reaching the target region was computed. This procedure was repeated 10 times for each initial condition, to compute a standard deviation for the target-hitting probability. Results are presented in Figure 6.

Note that since the three policies showed little qualitative difference, the three- and four-region policies were not investigated. Results indicate that for all of the initial conditions, the learned policy is likely to guide the system to the target region. In particular, the policy is very effective at initial conditions 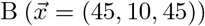 and 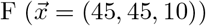, since from these conditions the optimal chemotherapy dose will not change along most likely paths to the target region, even if these paths slightly deviate from the path identified in Figure 5. Though this is also true for the initial condition 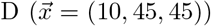, in practice it is more difficult to restore the *S* population using natural selection than it is to restore either of the resistant populations using chemotherapy. This phenomenon is explained in more detail in Section IV E. In contrast to these three initial conditions, any paths to the target region from the points 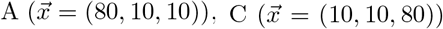, or 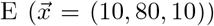 will require frequent switching of the chemotherapy dosage. This implies that the choice of action is, in a sense, less certain than in regions where a uniform dosage can be prescribed. Moreover, these points are simply further from the target region, and require longer paths to reach it. Hence, there are more opportunities for the state to deviate from the expected trajectory. It is also clear that the initial conditions C and E are most difficult for the learned policy to drive to the target region, likely because they begin with low populations of the *S* population, which again is difficult to recover from.

**FIG. 5.**
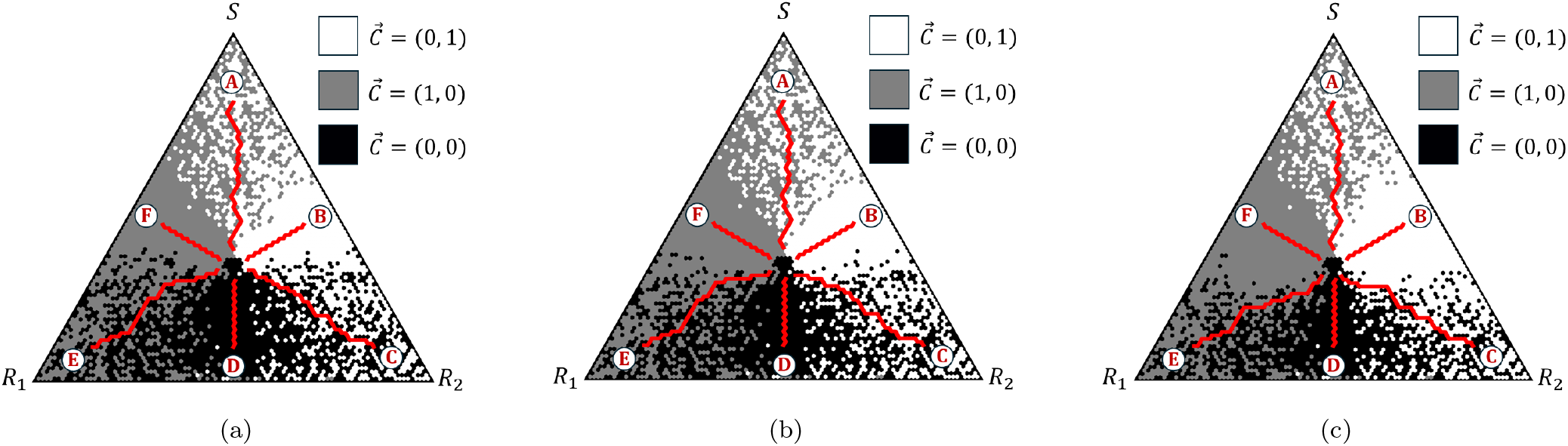
Most likely trajectories (red) with initial conditions chosen from the six different policy regions: *A* = (80, 10, 10), *B* = (45, 10, 45), *C* = (10, 10, 80), *D* = (10, 45, 45), *E* = (10, 80, 10), *F* = (45, 45, 10). (a) Two-region; (b) Three-region; (c) Four-region.

**FIG. 6.**
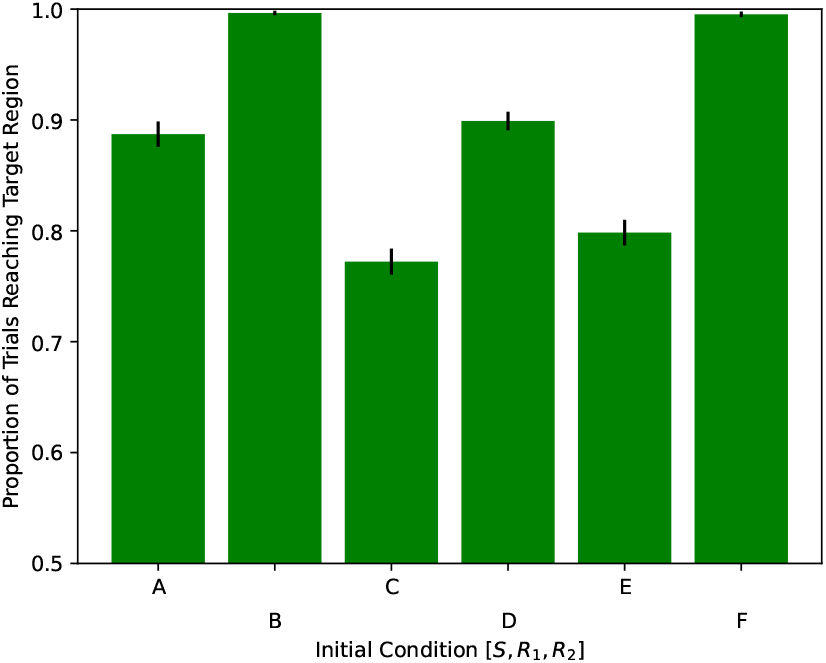
Probability of hitting the target region from each of the six initial conditions investigated in Figure 5, using the two-region policy.

### C. Robustness

A reasonable concern for these policies is their sensitivity to uncertainty in initial condition. The results presented in Figure 5 assume perfect knowledge of the cell subpopulation balance at the start of treatment. In this section, we quantify the ability for the trajectories to reach the target region when there is uncertainty in the initial state. Since no significant differences were identified between the three learned policies, we describe the results using only the two-region policy (shown in figure 2a).

Suppose that the true initial condition, 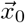, is not known, but can be estimated by some value 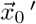. The objective of this analysis is to determine how well the chemotherapy policy *π* performs as the difference between 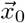 and 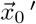 grows. To achieve this, a locally most probable trajectory is computed according to the procedure outlined in the previous section, using the initial condition 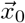. Concurrently, the corresponding chemotherapy schedule 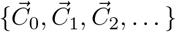 is computed. Then, *using the same chemotherapy schedule*, a new most likely path is computed from the estimated initial condition 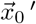 using Equation 8 to identify a new, adjusted locally most likely path.

Specifically, the state is initialized to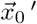 and from this state, the action 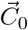 is taken. Transition probabilities to neighbors of 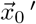 are computed, and the neighboring state with the greatest transition probability is denoted 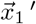. The process repeats by taking action 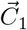 at 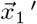 to identify the next most likely state 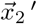, and then by taking action 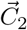 at 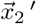 to identify 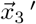 etc., until the action sequence 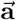 is exhausted. The adjusted locally most likely path is then taken to be the sequence 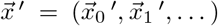.

Figure 7 shows six representative trajectories. Here, the action sequence is computed from the initial condition 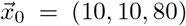, located towards the bottom-right of the simplex, and this action sequence applied to a range of different initial conditions 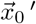 throughout the simplex.

**FIG. 7.**
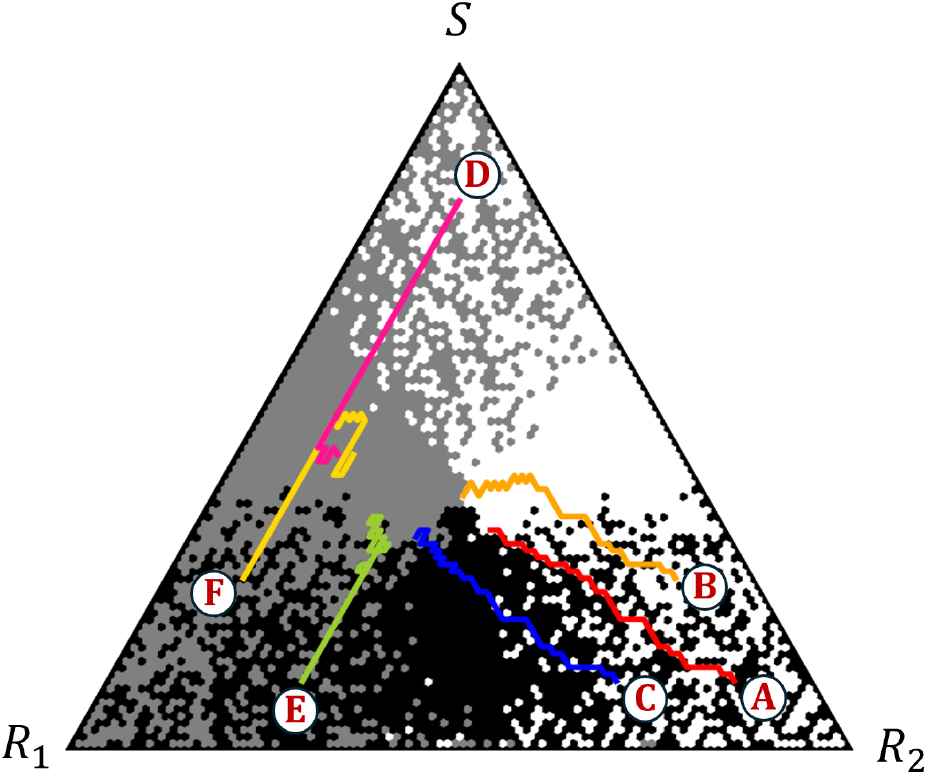
Locally most likely paths from each of six initial conditions when following the prescribed action sequence associated with the path from (A) 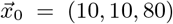. IC as follows: (A) 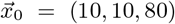, (B) 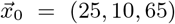, (C) 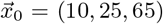, (D) 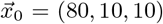, (E) 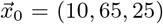, (F)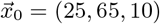.

Of course, the action sequence performs best when applied from initial conditions 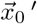 that are at or close to 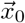. The path from 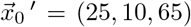 (orange) actually manages to reach to target region, while the path from 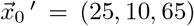 (blue) narrowly misses. In contrast, paths from completely different zones of the simplex miss the target completely, as expected.

Figure 8 quantifies robustness more systematically, where we plot the proportion of paths that reach our target vs. a measure of uncertainty in the initial condition. Each curve on this plot represents a different action sequence, computed from a different initial condition 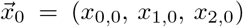. This action sequence is then applied to every other initial condition 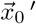 within a specified distance *d* of 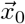, producing a series of adjusted trajectories. The proportion of these trajectories that reach the target region is observed and plotted against the distance *d*. Note that here, the distance between two states is taken to be the maximum difference in the size of any individual cell subpopulation between them, i.e.

**FIG. 8.**
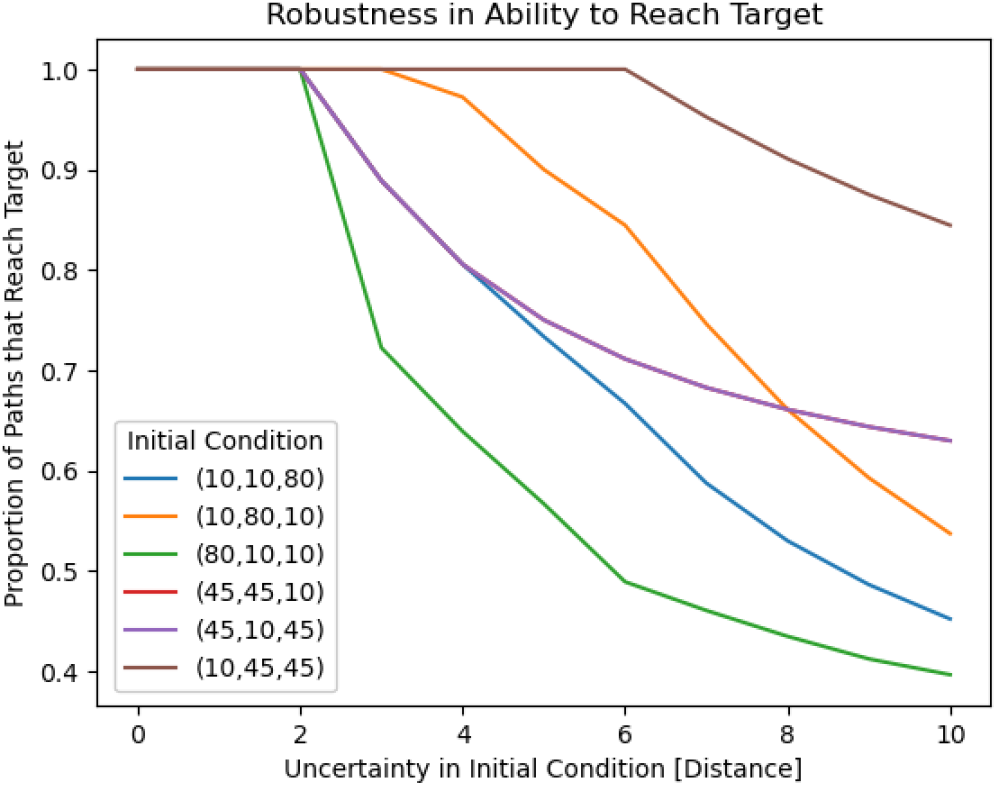
Proportion of paths reaching target region with initial conditions 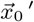 within the specified uncertainty range centered around 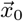 using that fixed action sequence.

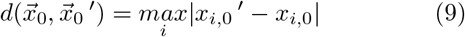

In general, it was found that small deviations (*d <* 3 in initial condition did not affect the ability of the extrapolated path to reach the target. It should be noted that the red curve corresponding to the IC 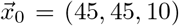 is hidden behind the purple curve corresponding to 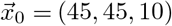. This is due to the symmetry between the *R*_1_ and *R*_2_ cell types (see Equation 1 and section II B), and the fact that the action sequences generated from these ICs uniformly prescribe the action 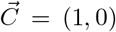 or 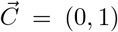 (see Figure 5). Interestingly, the policy is more robust in certain zones of the simplex than in others. At the initial condition 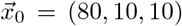 near the *S* corner of the simplex, the policy exhibits its least robust behavior (green line in Figure 8). In the opposing zone directly below the target, the policy appears most robust, as every trajectory starting within six units of the of the initial condition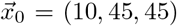 is able to reach the target.

This behavior is explained as follows. Near the *S* corner of the simplex, both of the resistant populations are at risk of extinction and the only actions prescribed by the chemotherapy policy are to apply *C*_1_ or apply *C*_2_, each of which applies selection pressure on one of the populations. Uncertainty (i.e. mismatches between the actions prescribed by the policy and the action sequence) made in this zone are more likely to result in extinction than similar uncertainties made elsewhere in the simplex. By contrast, in the zone below the target, the most probable trajectory from 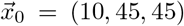 yields an action sequence where chemotherapy is never administered (see Figure 5). Mistakes in action are less likely here since the chemotherapy policy mostly prescribes 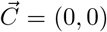 in this zone, and furthermore any mistakes that do occur are less likely to result in extinction.

Despite a clear discrepancy in robustness among the different ICs 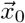, even in the worst case 40% of paths originating from within a 10-distance neighborhood of the IC from which action sequence was derived were able to reach the target region. This neighborhood of radius 10 corresponds to 270 unique states. By comparison, the target region itself contains only 42 states, with a maximum distance of 7 between any two points in this region (i.e. a radius of less than 4). Our general conclusion is that even with a relatively large degree of uncertainty in initial condition, a sizeable portion of most probable trajectories will still reach the target.

### D. Simplified policy

The learned chemotherapy policies presented in Figure 5 often prescribe alternating chemotherapy doses from step to step. These type of fine-scale switches in chemodosing is difficult to achieve in a clinical setting. Moreover, the analysis presented in Section IV C seemed to imply that policies prescribing uniform actions over large zones of the simplex were more robust when faced with uncertainty in measurements of the initial condition. In response to these observations, we investigated a simplified chemotherapy policy involving fewer action switches. Recall from Figure 4 that the learned policies can be divided into six zones. In the zones opposite each corner of the simplex, the policies largely prescribe a single action, while in the corner zones the policies prescribe a balance between a pair of actions. Figure 9 provides a simplified chemotherapy policy that capitalizes on this six-fold symmetry.

**FIG. 9.**
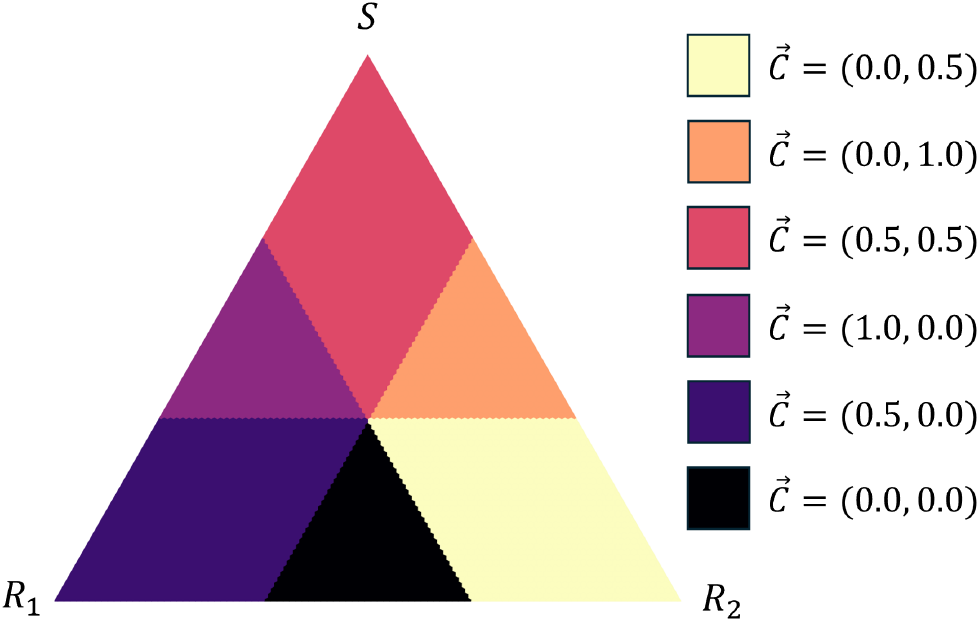
Simplified policies constructed by using the six-zone symmetry from figure 4. Uniform chemotherapeutic actions are assigned to each region as obtained from averaging the mixtures identified in figure 4.

**FIG. 10.**
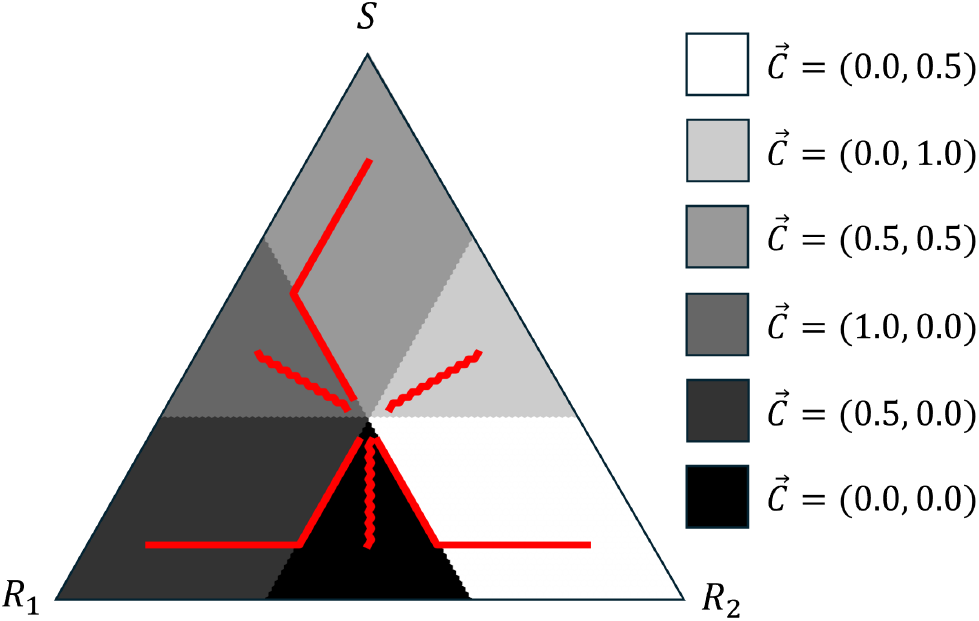
Locally most probable trajectories from each of the six zones, using the simplified policies shown in figure 9.

The six zones of the simplified policy are created by dividing the simplex along the *N/*3 axis of each sub-population, and a uniform action is prescribed in each zone. Notably, in the corner zones the prescribed action is a compromise between the two actions chosen by the learned policy. For example, in the zone closest to the *S* corner, the learned policy prescribes the actions 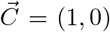 and 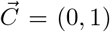 in an even ratio. In the simplified policy, this zone is therefore assigned the action 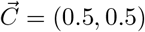.

The most probable trajectories were computed for the simplified policy, and are presented in Figure 10. All trajectories reached the target region, though paths with ICs near the corners of the simplex were found to approach the target quite differently from the paths shown in Figure 5. Specifically, the path with IC 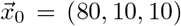, near the *S* corner, veered toward the *R*_1_ corner before turning toward the target. For the first step of this path, the *R*_1_ and *R*_2_ populations are perfectly symmetric and 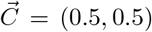. The path could proceed towards either the *R*_1_ or *R*_2_ corners with equal probability, and the *R*_1_ direction is chosen arbitrarily. However, on the next step 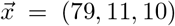 and the resistant populations become unbalanced. The species that was arbitrarily chosen for reproduction in the previous step becomes marginally favored in this one, and continues to be favored until the system reaches a state that prescribes a different chemotherapy dose. A similar phenomenon can be seen in the lower corners. For example, in the path with IC near the *R*_1_ corner, the *R*_2_ species is favored for reproduction over *S* from the first step, and as the *R*_2_ population grows it becomes increasingly favored until it reaches a state where the chemotherapy dose is switched.

In contrast, the learned policies frequently switch chemotherapy doses to alternately favor each of the species, and achieve paths that progress more directly towards the target. In a stochastic setting, moving directly toward the target is far safer as it immediately increases species diversity at every step and therefore mitiges the risk of species extinction through drift. Therefore, despite all of the constructed policy’s most probable trajectories eventually reaching the target, it is expected to be outperformed by the learned policies in numerical trials.

### E. Stochastic trials

In this section, we show the results from several numerical experiments associated with the learned chemotherapy policies. These simulations expose stochastic phenomena that would otherwise be difficult to identify by considering only the locally most likely trajectories. To introduce these simulations, Figure 11 shows two sample paths generated using the learned two-region chemotherapy policy, each from a different initial condition. The trajectories are generated using pseudo-random number generation to select, at each time step, one cell for reproduction and another for elimination.

**FIG. 11.**
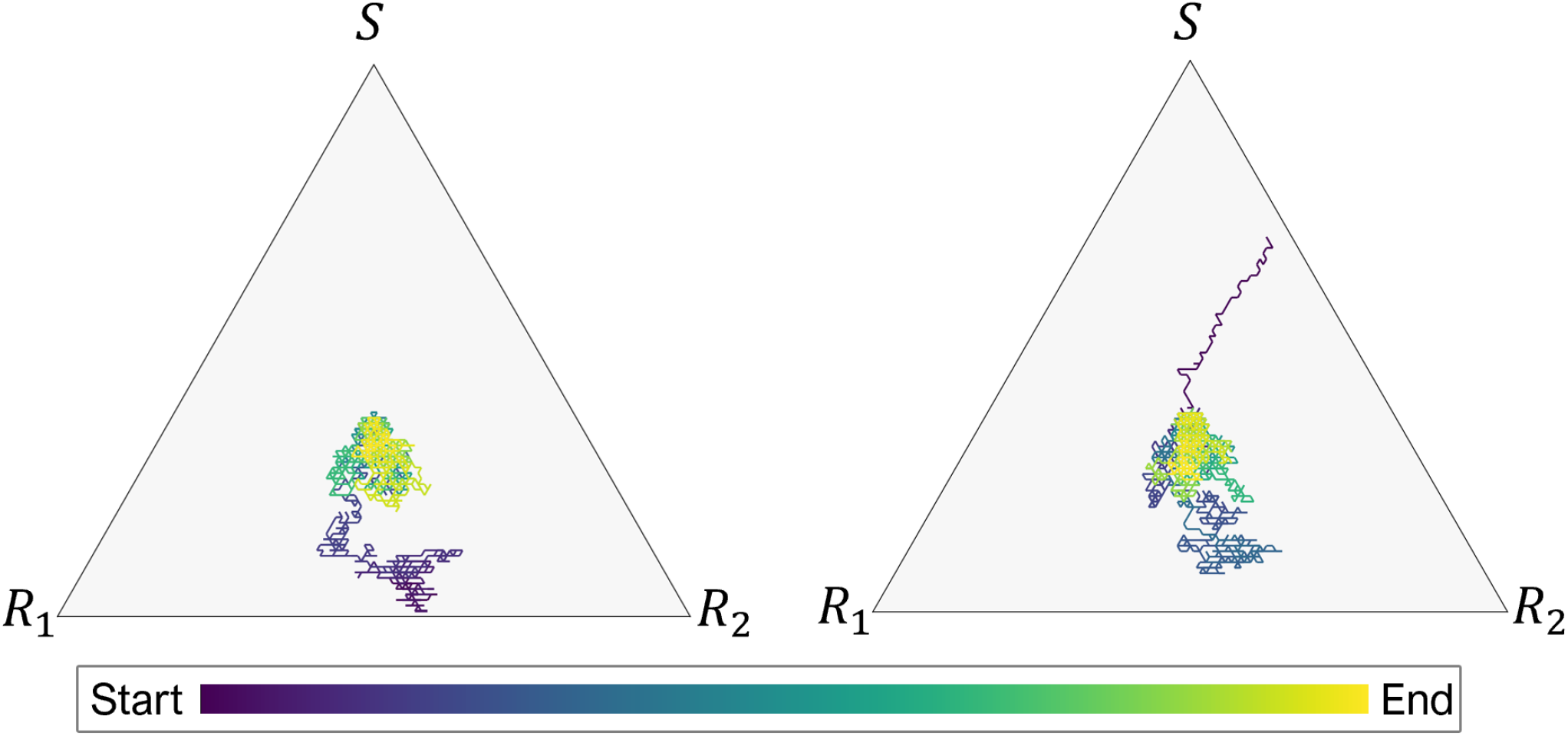
Two sample stochastic paths, with initial conditions 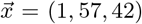 (left), and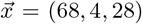 (right). Each path is colored by time: early time (dark blue); late time (yellow).

**FIG. 12.**
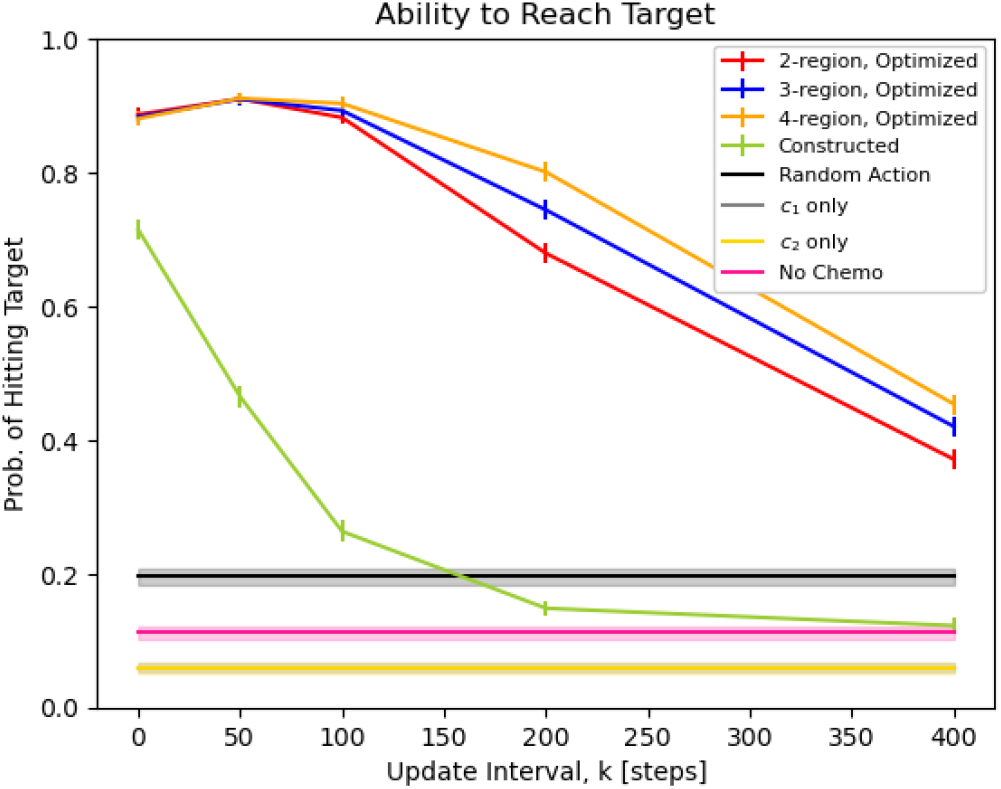
Probability of reaching target with incomplete information. Policy is updated every *k* steps (x-axis). Probability decreases as update interval increases. Also shown are sample benchmarks.

The paths in Figure 11 each contain 5000 steps. Both begin near an edge of the simplex (i.e. are in danger of extinction of one of the three subpopulations), and traverse the state space to reach the target. The first path meanders towards the target, but remains tightly within the target region upon arrival. The second path heads more directly towards the target, but later experiences some random downward drift before slowly correcting itself and returning to the target region. While these are only two stochastic realizations of a trajectory, they demonstrate a key behavior that is not captured by the analysis of the previous section.

Recall from Section III C that each of the three cell subpopulations is favored by exactly one of the allowable chemotherapy doses in the action space 𝔸. However the extent to which each cell type benefits under its favorable dose is not equal. To demonstrate this, consider the following three situations:

1. 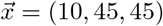, and action 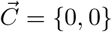 is chosen
2. 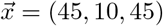, and action 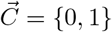 is chosen
3. 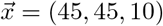, and action 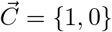 is chosen

In each case, one cell type is at risk of extinction and the chosen action is the one that favors the endangered cell type. The situation appears symmetric, however by computing the fitness vector 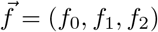 for each case (per the model presented in section II), it is straightforward to see that the ratio of the fitness of the favored cell type to that of the targeted cell types in each of the cases is (Case (i)) 1.33; (Case (ii)) 1.83; and (Case (iii)) 1.83. In Case (i), the endangered population is favored over its competitors by a slimmer margin than in Cases (ii) and (iii). It is still more likely for an *S* type cell to reproduce here than it is for either of the other species, but the difference in probabilities is less than that encountered in Cases (ii) and (iii). This means that in a stochastic setting, it is more difficult to drive the system state towards the *S* corner using the action 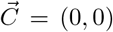 than to either of the other corners by using the actions 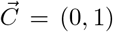 or 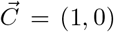. In other words, the effect of chemotherapy against sensitive cells is far stronger than the effect of natural selection against resistant cells. The consequence of this feature is that trajectories originating in the bottom half of the simplex are more prone to meander than those in the upper half, and random drift is more likely to result in extinction of the *S* population than either of the resistant populations.

### F. State-space estimates

Since each stochastic path includes the effect of random drift, a seperate element of uncertainty in the current state arises at each time-step where state-space postion measurements are not performed. In practice, this effect can be important since the exact state of the tumor can only be measured/estimated infrequently compared with the timescale on which the tumor cell balance changes. Therefore, the ability of the RI computed chemotherapy policy to perform well without perfect knowledge of the system state is investigated here.

To do this, we consider the disrepancy between what we call the true state of the tumor vs. the estimated state of the tumor. The true state of the system advances at each step according to the Markov process and choice of action, in the same manner as the paths presented in Figure 11. The estimated state is our best running estimate of the true state, which we use to choose actions at each time step. We assume that we know the initial state of the tumor with certainty, and can only update our exact knowledge of the state-vector every *k* steps thereafter. For *k* = 1, the true state and the estimated state coincide. For *k >* 1, the two states will diverge based on our imperfect knowledge. The larger the value of *k*, the more the paths should diverge.

Consider a state-space measurement taken at time *t* which reveals the true state 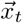. At that time, our estimated state is updated to match it. In between sequential state-space measurements, at the times *t*^*′*^ ∈ {*t*+ 1, *t*+ 2 … *t*+*k* − 1}, the true state is not known. To estimate it, the state is computed as the most probable trajectory with initial condition 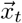, and total trajectory length *t*^*′*^. Actions can then be chosen based on this estimated state, until time *t*+*k* at which point the estimated state can once again be corrected to match the true state.

Experimental trials were performed to evaluate the ability of various chemotherapy policies to guide the system state towards the target under these assumptions using *k* as an uncertainty parameter. For each policy and update interval, 100 sets of 1000 numerical trials were performed, for a total of 100, 000 simulated trajectories. The initial condition for each trial was selected using pseudo-random number generation, with the only restrictions being that the initial condition is neither on an edge nor a vertex of the simplex, nor at the target region. Experiments were terminated upon reaching the target region (which counts as a success) or reaching an edge of the simplex (which counts as a failure). Within each set, the proportion of successful trials is computed. This proportion is then averaged across all 100 sets, and a sample standard deviation is computed as a measure of uncertainty. The results of this experiment are presented in Figure 12 where we plot the probability of hitting the target region vs. the uncertainty parameter *k*.

In addition to the constructed policy (Figure 9) and three optimized policies (Figure 4), for comparison purposes, Figure 12 also shows the performance of three additional policies: (1) *C*_1_ **only**, which takes the action 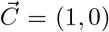 at all times; (2) **No chemo**, which takes the action 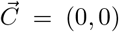 at all times; and (3) **Random action**, which chooses, at each time step, a random one of the three allowable actions *a* ∈ 𝔸. Since there is a direct symmetry between the species *R*_1_ and *R*_2_ and the chemotoxins *C*_1_ and *C*_2_, the policy *C*_2_ **only** is redundant and therefore excluded.

For small values of *k*, the three learned policies perform very similarly, with all policies successfully guiding the state towards the target in more than 90% of experimental trials. As *k* increases, the four-region policy outperforms the three-region policy, which in turn outperforms the two-region policy. The four-region reward structure and the policy it produces seems to be more resilient to uncertainty in the estimated state. But all three reward structures produce policies that yield a roughly 40% success rate at an update interval of 400 steps, which is long enough to traverse from one side of the simplex to another many times over.

By contrast, the simplified policy does not perform nearly as favorably. For *k* = 0 (i.e. with perfect knowledge of the system state at all times), the simplied policy achieves a moderately high rate of success, but sees a sharp drop-off in performance as *k* increases, eventually performing worse than the “random action” benchmark at intervals exceeding 100 steps. Our conclusion, in general terms, is that the simplified policy can be reasonably effective, but is generally not robust.

## V. CONCLUSION

Reinforcement learning/machine learning algorithms have great potential to sort through many possible drug combinations and temporal schedules for individual patients before decisions are made to determine which therapies are optimal. The larger the population, the finer the deliniations in the population subtypes, and the more drug combinations and timing involved all provide challenges given the generally slow convergence of the algorithms (i.e. they suffer from the curse of dimensionality). But these same complications also plague any alternative decision making process. With better mathematical models (which can speed up convergence), and more sophisticated real-time data assimilation (which can tailor models to individual patients), algorithmic methods offer great potential for designing complex drug-combinations and dose-timing protocols for eventual clinical utility.

## ACKNOWLEDGMENTS

We gratefully acknowledge support from the Army Research Office MURI Award #W911NF1910269 (2019-2024).

## Notes

### Competing Interest Statement

The authors have declared no competing interest.

